# Age-Related Differences in Carbohydrate and Fat Oxidation Over 24 Hours and During Daily Activities in Humans

**DOI:** 10.64898/2026.09.24.754171

**Authors:** Zoe H. Smith, Christopher M.T. Hayden, Luke R. Arieta, Michael A. Busa, Jane A. Kent

## Abstract

The effects of older age on the oxidation of energy substrates (carbohydrates, fats) over 24 hr and during activities of daily living (ADL) may have substantial implications for metabolic health. To examine this issue, we evaluated age-related differences in substrate oxidation for 24 hr and periods of sleep, ADLs, and a 30-min treadmill walk (30MTW, 1.3m·s^-1^) in 17 young (29 ± 5 yr, mean ± SD, 6F) and 16 older (72 ± 5, 8F) healthy active adults. Oxygen consumption (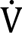O_2_) and carbon dioxide production were measured using a 32,500 L room calorimeter to quantify substrate oxidation, which was normalized to body mass (kJ·kg^-1^) and total energy expenditure (%). Separately, 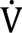O_2peak_ was measured during an incremental treadmill test to exhaustion. 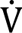O_2peak_ (ml·kg^-1^·min^-1^) was lower in older adults compared with young adults. Older adults oxidized fewer carbohydrates than young over 24 hr (kJ·kg^-1^, %), sleep (kJ·kg^-1^), and ADLs (kJ·kg^-1^, %); and fat oxidation was greater in older than young over 24 hr (kJ·kg^-1^, %) and ADLs (kJ·kg^-1^, %). Carbohydrate oxidation did not differ by group during the 30MTW (kJ·kg^-1^, %), but fat oxidation was greater in older (kJ·kg^-1^) despite their walking at a greater relative intensity. These results indicate that older adults derived oxidative energy differently than young over 24 hr and during daily tasks. The lower carbohydrate and greater fat contribution to daily energy requirements may be meaningful in the development of metabolic disease in older adults.

**Key Points:**

- The extent of age-related differences in energy substrate (carbohydrate and fat) oxidation remains unclear, particularly over long periods of time and during daily tasks.
- We evaluated substrate oxidation in healthy young and older males and females over 24 hours that included sleep, activities of daily living (ADLs) and a 30-minute treadmill walk at 1.3 m·s^-1^.
- The older group oxidized less carbohydrate over 24 hours, sleep, and ADLs; and more fat over 24 hours and during ADLs than young.
- Despite walking at a greater relative intensity, carbohydrate oxidation was not different and fat oxidation was greater in older compared with young.
- These results show distinct differences in substrate oxidation, mainly in the form of lower carbohydrate and greater fat oxidation, in older compared with younger adults that are evident over 24 hours, sleep and during daily physical tasks. The implications for metabolic health in older adults remain to be determined.

## INTRODUCTION

The direct energy (ATP) used by the body to fuel biochemical processes, such as the fundamental interactions between myosin and actin involved in movement, is produced by the breakdown of energy-containing substrates derived from food (carbohydrate, fat, and protein). Energy requirements at rest and during low-to-moderate intensity exercise are met primarily through the oxidation of carbohydrate and fat. Indirect calorimetry, which measures oxygen consumption (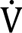O_2_) and carbon dioxide (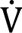CO_2_) production by gas exchange, is used to calculate total energy expenditure (kJ), as well as carbohydrate and fat oxidation via the respiratory exchange ratio (RER; 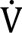CO_2_/ 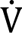O_2_) or established equations (kJ; Brouwer, 1957). However, measures of gas exchange are typically made with a mask or mouthpiece connected by a hose to a gas analyzer, which generally limits both the movement of the individual to a stationary device (e.g., chair, bike, treadmill) and the time over which the measures can be comfortably obtained. Whole-room indirect calorimetry (Chen et al., 2020) can overcome these limitations as gas exchange, and thus substrate oxidation, can be recorded without restricting movement and over a full day while a person is in the room. Most researchers who have taken advantage of room calorimetry have done so to investigate nutritional or exercise interventions, metabolic disease, and obesity (Carnero et al., 2021; Melanson et al., 2005, 2002; Rynders et al., 2018). In contrast, few have used room calorimetry to explore the question of whether young and older adults use energy substrates differently over a day. The implications of altered substrate selection in older age may include metabolic inflexibility (Kelley et al., 1999), obesity (Goodpaster et al., 2002), and insulin resistance (Petersen et al., 2015).

The results of the few studies that have used room calorimetry to investigate age-related differences in substrate oxidation are equivocal (Davy et al., 2001; Levadoux et al., 2001; Melanson et al., 2007). Davy et al., (2001) concluded that there were no age-related differences in 24-hr substrate oxidation (measured as % total energy expenditure), while others have reported lower (Melanson et al., 2007) or greater (Levadoux et al., 2001) carbohydrate oxidation in older than young adults. Given the conflicting information to date, a consensus about how age may influence substrate oxidation over 24 hours has not been established. Additionally, there is little information available regarding age-related differences in substrate oxidation during sub-periods within the day, such as sleep and activities of daily living (ADL). Thus, whether and how substrate oxidation during daily tasks influences 24-hour measures among young and older adults is unclear.

The purpose of this study was to determine whether there were age-related differences in substrate oxidation over a 24-hr period that included sleep, a period of activities of daily living (ADLs), and a 30-minute treadmill walk at 1.3 m·s^-1^ (30MTW, Foulis et al., 2017). Due to the lower aerobic capacity (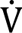O_2peak_) of older than young adults (Kent-Braun and Ng, 2000; Ogawa et al., 1992), and the well-accepted dominance of carbohydrate for energy production at higher relative exercise intensities, we hypothesized that older adults would oxidize more carbohydrate and less fat compared with young during 1) the ADLs and 30MTW, and 2) over the full 24-hr period. Our results highlight distinct age-related differences in substrate oxidation during daily tasks and over 24 hours that should be considered in future work aimed at understanding the development of adverse metabolic health outcomes or disease (i.e., obesity, insulin resistance) in older age.

## METHODS

Following written informed consent, collection of descriptive data (e.g., height, body composition), and familiarization with study procedures, young and older participants underwent a peak oxygen consumption (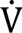O_2peak_) test and a 24-hr room calorimeter visit. The 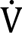O_2peak_ and 24-hr room calorimeter visit were separated by at least 72 hours.

### Ethical Approval

This study was submitted to and approved by the University Institutional Review Board and conducted in accordance with the Declaration of Helsinki., with the exception of pre-registration in a database. All participants provided written informed consent prior to any data collection. Portions of the data from the 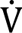O_2peak_ test in the young males have addressed different scientific questions and are reported elsewhere (Hayden et al., 2025).

### Participants

Sedentary-to-moderately active young (25-40 yr) and older (65-80 yr) males and females were recruited from the University and surrounding communities. Inclusion criteria were: free from known disease and relatively weight stable (<5% change in preceding 6 months). Exclusion criteria were: neurological, neuromuscular, cardiac, pulmonary or metabolic disease; taking medication that would affect neuromuscular or metabolic function; rheumatoid arthritis, uncontrolled hypertension (blood pressure >140/80), smoking, history of symptoms upon physical exertion, BMI <18 or >35 kg·m^-2^, or surgery in the preceding 6 months that would impact the ability to perform physical activity. A signed health care provider’s clearance form was required for all older adults prior to participating in the study.

### Descriptive Data and Familiarization

Height (m) was measured for all participants and blood pressure (mmHg) was obtained for all older adults to screen for hypertension. A dual-energy X-ray absorptiometry scan was completed (Lunar iDXA, General Electric, Chicago IL) to obtain measurements of body mass (kg), lean mass (% total), and fat mass (% total). A Short Physical Performance Battery (SPPB, Guralnik et al., 1994) consisting of walking, chair rise, and balance tests, was completed by all older participants to characterize physical function in this group. Next, participants were familiarized with walking on the same treadmill to be used during the 24-hr room calorimeter visit (described in detail below).

Each participant was instructed to wear an ActiGraph GT3x+ accelerometer (Ametris, Pensacola, FL) on their right hip and to fill out a physical activity log for 7 days. The participant was asked to record all activities in the log, as well as when the accelerometer was put on and taken off each day. These data were used to characterize free-living physical activity in our groups. The following variables were determined over the 7 days using ActiLife v6.13 software (Ametris, Pensacola, FL): average daily physical activity counts (vector magnitude counts·d^-1^·1000^-1^), weekly minutes of moderate-to-vigorous physical activity (min·week^-1^), and average daily steps (steps·day^-1^). Minutes of moderate-to-vigorous physical activity were determined using established cut points (Troiano et al., 2008). Participants were also asked to fill out a diet log for three days, which consisted of each participant’s written record and photos of all food and beverages consumed for these days. This diet log was used to design the diet for each participant’s room calorimetry visit (described below), with the goal of closely mimicking their typical diet.

### Peak Oxygen Consumption (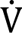O_2peak_) Visit

All participants were asked to refrain from alcohol consumption or heavy exercise the day prior to, and to avoid exercise and caffeine the morning of, this visit. Participants were instructed to eat breakfast as usual, and to consume a standardized meal bar (1,808 kJ, PROBAR LLC, Salt Lake City, UT) between 10:30 and 11:30 am. A modified Bruce ramp protocol on a treadmill (Quinton TM 55, Davis Medical Electronics, Inc., Vista, CA) was performed between 1:00 and 3:00 pm to determine 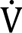O_2peak_ (Kaminsky and Whaley, 1998). The test began at a speed of 0.76 m·s^-1^ and 0% grade. Grade was increased by 1.2-1.3 % every 20 s for the first 3 min of the test, after which both speed and grade were increased. Verbal encouragement was given to the participant throughout the test. Oxygen consumption and 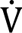CO_2_ were measured continuously using a Parvo Medics TrueOne 2400 metabolic cart (Parvo Medics, Salt Lake City, UT). The test was terminated either when the participant reached volitational fatigue or when the study staff determined they could not safely continue.

A rating of perceived exertion was recorded every minute of the test using a modified Borg scale from 0 to 10, with 10 being maximal exertion (Borg, 1982). For the older adults, a 12-lead electrocardiogram was positioned and monitored before, during, and following the test by a clinical exercise physiologist, and blood pressure was taken manually every 2 min. Data were exported from the metabolic cart with a time resolution of ∼5 s. A 30-s moving average was applied to the data using a custom MatLab script (V2020b, MathWorks, Natick, MA). The moving average was then used to obtain peak oxygen consumption (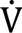O_2peak_; L·min^-1^) and the respiratory exchange ratio (RER) at 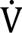O_2peak_. Peak 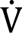O_2_ was normalized to body mass (kg) for each participant.

### Blood Draws and Analyses

Blood draws were performed for young female participants either the day before or during the 24-hr room calorimeter visit to determine reproductive hormone concentrations, as some (Hackney et al., 1994), but not all (D’Souza et al., 2023), studies suggest that these hormones may influence substrate oxidation. ELISA assay kits were used to determine serum concentrations of progesterone (ng·mL^-1^), estradiol (pg·mL^-1^), follicle-stimulating hormone (mIU·mL^-1^), and luteinizing hormone (mIU·mL^-1^).

### 24-hr Room Calorimeter Visit

At least 72 hours following the first visit, participants arrived at the laboratory at 6:45 pm prepared to stay in the room calorimeter (MEI Research, Ltd, Minneapolis, Minnesota) until 6:00 pm the next day. Young female participants took part in this study visit approximately 5-10 days after the start of their last menstrual cycle, except for one young female who did not have a regular menstrual cycle due to hormonal contraceptive use. The room calorimeter was 4.0 x 3.4 x 2.6 m in size and equipped with a bed, desk, chair, treadmill, toilet, sink, television, window with a view of the campus, window into the laboratory space from which the researcher monitored all testing, and two wall ports through which to pass food and supplies. The temperature of the room was set to 22° C for all 24-hr visits. Prior to entering the chamber, body mass (kg) was measured for all participants, and blood pressure was obtained for older adults. A research team member was present continuously during the stay and communicated with the participant by walkie-talkie. The schedule of activities for this visit is shown in Table 1. Extensive calibration, validation, and analytical procedures were conducted prior to this study (Hayden et al., 2026).

**Table 1.** Room calorimeter schedule.

| Time | Activity |
| --- | --- |
| 7:00 PM | Enter room calorimeter |
| 7:15 PM | Dinner (individualized) |
| 12:00 AM | Participant must be in bed |
| 8:00 AM | Participant must be out of bed |
| 8:15 AM | Activities of daily living (ADLs) |
| 9:15 AM | Breakfast (individualized) |
| 11:30 AM | Lunch (1,808 kJ standard meal bar) |
| 12:00 PM | Must have completed 1,200 steps |
| 2:00 PM | 30-min treadmill walk (30MTW) |
| 2:30 PM | Seated 2-hour recovery |
| 6:00 PM | Exit room calorimeter |

Physical activity was measured throughout the 24-hr visit using an ActivPal (PAL Technologies LTD, Scotland, UK) device placed with an adhesive film to the participant’s right thigh, approximately halfway between the knee and the hip. The ActivPal rather than the ActiGraph device was used while in the chamber because the ActivPal can detect subtle changes in movement including position changes from lying or sitting to standing (Grant et al., 2006), and it was expected that less overall movement would be performed while in the chamber compared with free-living due to space constraints. These data were used to quantify time spent sitting or lying down, standing, and stepping; as well as overall physical activity for each group while in the room calorimeter.

Each meal was provided by the research staff through the wall port. Dinner and breakfast were matched for caloric and macronutrient intake according to each participant’s usual diet, based on their 3-day diet log. For lunch, a standard meal bar (1,808 kJ, PROBAR LLC, Salt Lake City, UT) was consumed by all participants approximately 2.5 hours prior to the 30MTW. The caloric and macronutrient content of the food and beverages consumed during the 24-hr visit was calculated based on nutrition labels, or using a nutrition application (MyFitnessPal, Francisco Partners, San Francisco, CA) when a nutrition label was not available.

All ADLs were performed before breakfast and consisted of four 10-min bouts of activity, each separated by 5 min of sitting. The activities included making the bed, folding laundry while standing, vacuuming, and washing dishes, and were done in this order by all participants. Each participant was guided through the activities by the research staff. All participants were then encouraged to accumulate 1,200 steps (including steps during ADLs) before noon to minimize sedentary behavior during the 24-hr stay.

The 30MTW took place ∼2.5 hours after lunch. Thirty minutes prior to the treadmill walk, the participants were asked to sit in a chair for 20 min and then to stand on the treadmill (Valiant 2, Lode B.V., Groningen, Netherlands) for 10 min to obtain baseline measurements. The treadmill was controlled by the research staff from outside the room. Once started, the treadmill increased in speed to 1.3 m·s^-1^, which was maintained by all participants throughout the test, apart from one older female who walked at a speed of 0.9 m·s^-1^. Treadmill grade began at 0% and was increased to 3% for 1 min at minutes 7, 17, and 27, to mimic real-life activities such as walking up a hill or stairs, referred to as “challenge periods” (Foulis et al., 2017). Participants were instructed not to grip the treadmill but were allowed to place two fingers on the side rails to help with balance, if needed. When the treadmill stopped at the end of the 30 min, participants stepped off immediately and returned to a seated position.

Oxygen and carbon dioxide gas analyzers (Siemens, Munich, Germany) measured the gases in voltages as the air exited the chamber, with sampling every 15 s. Details regarding the calibration and analysis of these data are provided elsewhere (Hayden et al., 2026). The oxygen sensor was paramagnetic, and the carbon dioxide sensor was infrared. Raw voltages were converted into concentrations in CalRQ (MEI Research, Ltd, Minneapolis, Minnesota). Oxygen values were corrected with a 12-point calibration fit with a linear equation and the carbon dioxide values were corrected based on an 11-point calibration fit with a third-order polynomial. A 0.0025 Hz Butterworth lowpass filter was applied before applying a 4-min centered derivative to the data. The 4-min centered derivative calculated the rate of change of the center point of data, taking into account the data 2 min before and after. The data were then processed using a custom MatLab script. The first hour and the final 15 min of the stay were discarded to ensure all metabolic data were a result of the participants’ 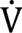O_2_ and not an artifact of fluctuations that can occur when a person first enters the room, or when the door is opened for them to egress, leaving a data collection period of ∼21-22 hr. Data were extrapolated to cover a full 24-hour period using an average of the entire dataset. Sleep was determined as the lowest 3 hours of energy expenditure from the time the participant went to bed until they got up in the morning. The time period for the ADLs was recorded by the investigator as the start to the end of the activities. The ADL time period started when the participant stood up for the first activity and ended after 5 min of seated rest following the final activity. The 30MTW data were selected from when the treadmill started to when it stopped, which was recorded alongside the metabolic data in the CalRQ file.

Variables directly recorded from CalRQ during each time period of interest were: 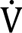O_2_ (L·d^-1^), 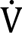CO_2_ (L·d^-1^), and energy expenditure (kJ·d^-1^). The RER was calculated using the sum of 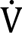O_2_ and 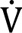CO_2_ during each time-period of interest, as recommended by Chen et al., (2020) to reduce the influence of noise on the measurement. The RER during the 30MTW was calculated from the sum of 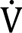O_2_ and 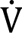CO_2_ from minutes 20 to 27, to avoid transients due to the onset of exercise and the challenge periods. Carbohydrate oxidation (g, Equation 1) and fat oxidation (g, Equation 2) were calculated as follows (Brouwer, 1957):

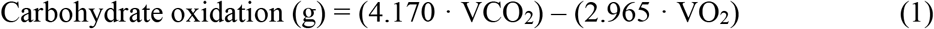

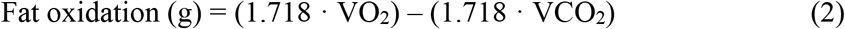

Protein oxidation was assumed to be negligible. Using 24-hr protein oxidation values for young and older adults from two studies in the literature (Levadoux et al., 2001; Melanson et al., 2007), a separate analysis was performed to determine whether, and by how much, including protein oxidation would influence any age-related comparisons of carbohydrate and fat oxidation over the 24-hr period in the current study. For this analysis, the following equations were used (Equations 3 and 4):

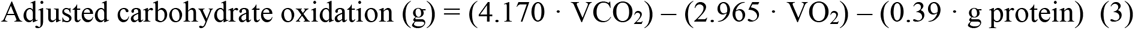

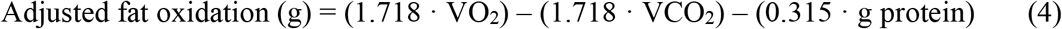

Protein oxidation values of 86 and 96 g for young and older adults from Melanson et al., (2007), as well as values of 85, 65, 92, and 62 g for young males and females and older males and females, respectively, from Levadoux et al. (2001) were used in this analysis. The protein oxidation results from Davy et al., (2001) were not used in this analysis as grams oxidized per day were not reported.

Total carbohydrate and fat oxidation were converted from grams to kilocalories (kcals) with a conversion factor of 4.182 kcal·g^-1^ for carbohydrate, assuming glycogen as the primary carbohydrate substrate, and 9.461 kcals·g^-1^ for fat (Brouwer, 1957; Lusk, 1917). Carbohydrate and fat oxidation were then converted from kcals to kilojoules (kJ) with a conversion factor of 4.186 kJ·kcal^-1^. Energy expenditure, carbohydrate oxidation, and fat oxidation were normalized to total mass of the participant, as recommended by consensus (Chen et al., 2020). Energy balance (kJ) was calculated as the difference in energy expended and consumed over the 24-hr visit.

To explore carbohydrate and fat oxidation in reference to the “crossover” concept of substrate use (Brooks and Mercier, 1994), plots were created for each substrate as a % of total energy expenditure *vs.* 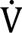O_2_ and %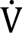O_2peak_, using the ADL and 30MTW data. Thus, each participant contributed 4 data points (mean carbohydrate and fat during ADLs and 30MTW) and linear fits were applied to the carbohydrate and fat data for each group, separately, to examine potential age-related differences in the crossover from fat to carbohydrate oxidation with increasing exercise intensity.

### Statistical Analyses

All statistical analyses were conducted using R Studio Statistical Software (Version 4.4.2, Posit team 2024). The Shapiro-Wilk test was performed first to determine whether the data were normally distributed. Two-tailed, independent student’s t-tests were performed to detect differences in group means for: descriptive characteristics; physical activity during the 24-hr visit; macronutrient and caloric intake (kJ) during breakfast and dinner; and 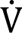O_2_, % 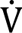O_2peak,_ 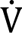CO_2_, energy expenditure, RER, fat oxidation, and carbohydrate oxidation for sleep, ADLs and the 30MTW (Hypothesis 1), as well as the 24-hr visit (Hypothesis 2). A nonparametric, Wilcoxon rank-sum test was used to detect group differences in energy expenditure during ADLs as well as time spent standing and stepping in the chamber, as these variables were not normally distributed. A p-value ≤ 0.05 was considered significant. Data are reported as mean ± SD, and precise p-values and the 95% confidence interval (CI) for differences between means are provided.

## RESULTS

### Descriptive Characteristics

Group characteristics are shown in Table 2. Seventeen young (6 female) and 16 older (8 female) adults participated in this study. There were no differences between the groups in height, body mass, lean mass, or fat mass. Physical activity data were extrapolated from 6 to 7 days for 1 young female, 1 older male, and 1 older female due to one day being excluded from analysis. No differences by age were detected for daily physical activity, weekly minutes of moderate-to-vigorous physical activity, or steps per day. The SPPB score for the older adults ranged from 9-12 (median = 11). No participants reported following a high- or low-fat or carbohydrate diet.

**Table 2.** Group characteristics.

|  | Young (n=17) | Older (n=16) | p value | 95% CI |
| --- | --- | --- | --- | --- |
| Age (yr) | 29 ± 5 | 72 ± 5 | - | - |
| Female (%) | 35 | 50 | - | - |
| Height (m) | 1.73 ± 0.11 | 1.71 ± 0.11 | 0.461 | -0.05, 0.11 |
| Body mass (kg) | 79 ± 16 | 71 ± 10 | 0.084 | -1, 17 |
| Lean mass (%) | 71 ± 9 | 67 ± 10 | 0.232 | -3, 11 |
| Fat mass (%) | 25 ± 10 | 30 ± 11 | 0.229 | -12, 3 |
| Physical activity (ct·d<sup>-1</sup>/1000) | 485 ± 215 | 500 ± 184 | 0.830 | -157, 127 |
| MVPA (min·week<sup>-1</sup>) | 353 ± 188 | 367 ± 195 | 0.841 | -150, 123 |
| Steps per day | 6,851 ± 3,114 | 8,017 ± 3,868 | 0.350 | -3,676, 1,345 |
| $\dot{V}O_{2peak}$ (ml·kg<sup>-1</sup>·min<sup>-1</sup>) | 42 ± 8 | 28 ± 7 | <0.001 | 9, 20 |
| RER at $\dot{V}O_2$ peak | 1.15 ± 0.07 | 1.07 ± 0.06 | 0.004 | 0.03, 0.12 |

Data are mean ± SD; ct, counts; MVPA, minutes of moderate-to-vigorous physical activity;
$\dot{V}O_{2peak}$ , peak oxygen consumption; CI, confidence interval for difference in means; RER,
respiratory exchange ratio; n=13 for $\dot{V}O_{2peak}$ measurements from older group.

### Peak Oxygen Consumption (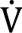O_2peak_)

Average 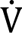O_2peak_ data for the young and older groups are shown in Table 2. As expected, 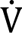O_2peak_ of the older group was lower than the young group, as was RER at 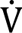O_2peak_. Two older males were unable to complete the 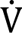O_2peak_ test due to the development of ECG irregularities, and one older female terminated the test early due to a high perceived exertion, leaving n=13 in the older group for this measurement and any measurements that were normalized to 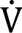O_2peak_.

### Blood Draw Outcomes

Average [progesterone] was 0.53 ± 0.29 ng·mL^-1^ (range: 0.26 to 0.94) in the young female participants, while [estradiol] was 99.0 ± 178.4 pg·mL^-1^ (range: 12.2 to 462.3). There were two participants on hormonal contraception, one of which had a normal menstrual cycle while the other did not. The participant who did not have a normal menstrual cycle was not limited to being in the room calorimeter during their follicular phase, as we were unable to determine when this occurred, and was an outlier with a high [estradiol] (462.3 pg·mL^-1^). The range in [estradiol] for the remaining participants (n=5) was 12.2 to 42.9 pg·mL^-1^. [Follicle-stimulating hormone] and [luteinizing hormone] were 5.4 ± 1.9 mIU·mL^-1^ (range: 2.4 to 7.4) and 9.0 ± 5.2 (3.8 to 16.1).

### Diet and Physical Activity During 24-hr Stay

There was no difference by age in the macronutrient and caloric intake of breakfast (Young: 1,472 ± 553 kJ CHO, 994 ± 442 kJ fat, 544 ± 255 kJ protein, 3,010 ± 926 kJ total; Older: 1,295 ± 567 CHO, 766 ± 580 fat, 405 ± 193 protein, 2,466 ± 899 total; p ≥ 0.086) or dinner (Young: 1,686 ± 792 CHO, 1,122 ± 645 fat, 672 ± 271 protein, 3,480 ± 1,424 total; Older: 1,366 ± 392 CHO, 1,111 ± 420 fat, 696 ± 289 protein, 3,173 ± 717 total; p ≥ 0.152).

The time spent sitting or lying down (Young: 19.7 ± 0.7 hr; Older: 19.5 ± 1.0; p = 0.490), standing (Young: 3.2 ± 0.6; Older: 3.6 ± 0.9; p=0.179), or stepping (Young: 1.1 ± 0.3; Older: 1.2 ± 0.3; p = 0.313) was not different by age group. For one young female and two older males, the time spent in each of these activities during the room calorimeter stay was not recorded due to device error, resulting in n = 16 for young and n = 14 for older for these measures.

### 24-hour Outcomes

Representative metabolic data from one younger and one older female spanning the 24-hr visit are shown in Figure 1. Oxygen consumption, VCO_2_, energy expenditure, and RER over 24 hours are given in Table 3. Although 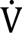O_2_ (summed and normalized to body mass) did not differ between groups, when expressed as a percentage of 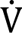O_2peak_, 24-hr 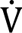O_2_ was greater in older compared with young. VCO_2_ was lower in the older than in the young. As a result, RER averaged over the 24 hours was lower in the older group compared with the young. Total and body-weight normalized energy expenditure were not different between groups.

**Figure 1.**
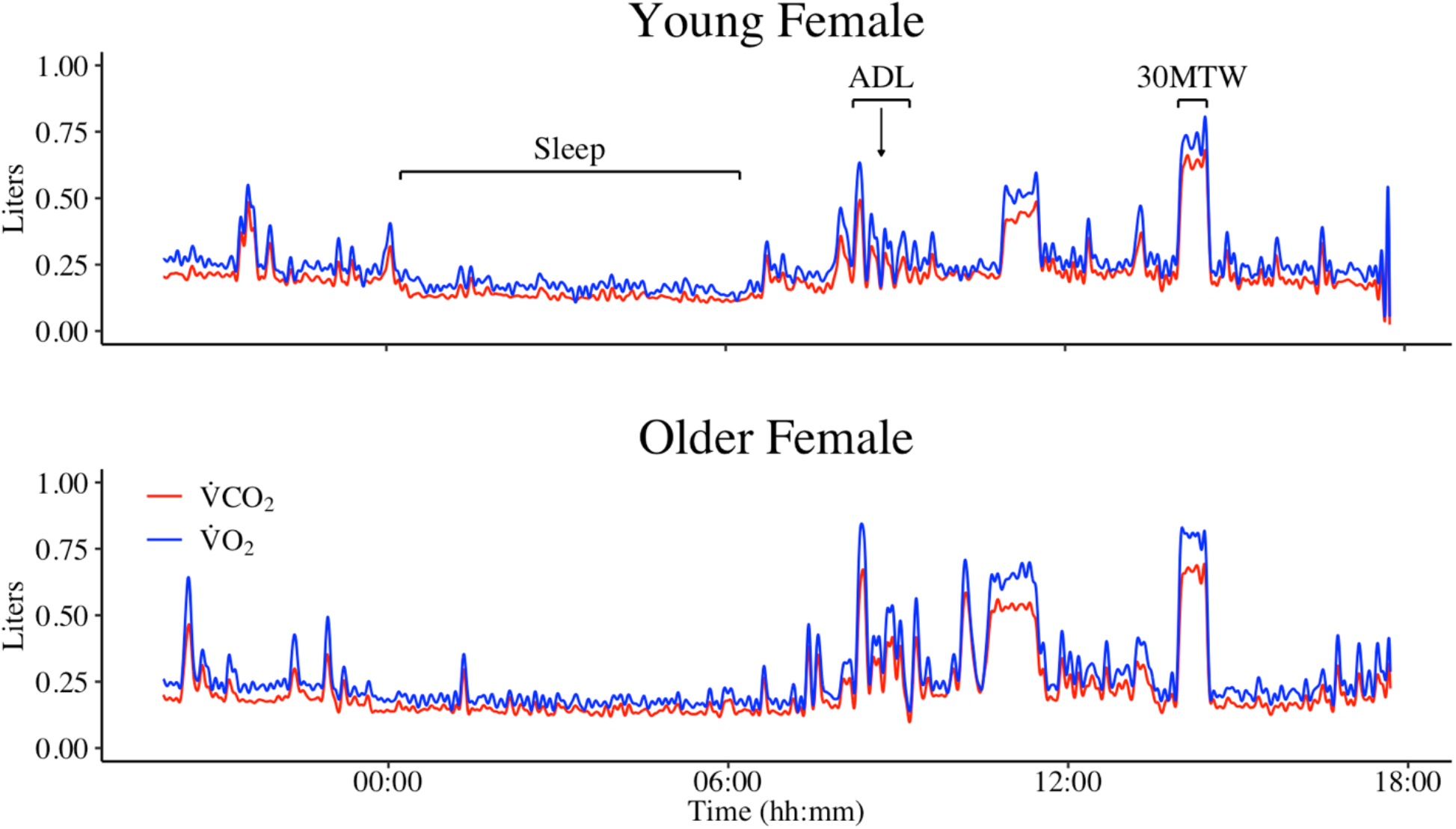
Representative gas exchange data from one young (top) and one older (bottom) female. Oxygen consumption (VO_2_, blue line) and carbon dioxide production (VCO_2_, red line) throughout the 24-hr room calorimeter stay are shown. Time periods of interest are identified in the top figure by brackets. The time between the ADLs and 30MTW for both participants included a period of self-paced treadmill walking, which aided in reaching the goal of 1,200 steps by noon.

**Table 3.** Metabolic variables for 24-hour period.

|  | Young (n=17) | Older (n=16) | p value | 95% CI |
| --- | --- | --- | --- | --- |
| Total VO<sub>2</sub> (L) | 490 ± 104 | 436 ± 75 | 0.095 | -10, 118 |
| VO<sub>2</sub> (ml·kg<sup>-1</sup>·min<sup>-1</sup>) | 4.4 ± 0.5 | 4.4 ± 0.5 | 0.869 | -0.4, 0.4 |
| VO<sub>2</sub> (% peak) | 11 ± 2 | 17 ± 3 | <0.001 | -8, -4 |
| Total VCO<sub>2</sub> (L) | 410 ± 88 | 355 ± 59 | 0.042 | 2, 108 |
| Energy Expenditure (kJ) | 9,895 ± 2,095 | 8,760 ± 1,490 | 0.082 | -153, 2,424 |
| Energy Expenditure (kJ·kg<sup>-1</sup>) | 126 ± 15 | 125 ± 16 | 0.762 | -9, 13 |
| RER | 0.84 ± 0.02 | 0.81 ± 0.02 | 0.003 | 0.01, 0.04 |

Data are mean ± SD; VO<sub>2</sub>, oxygen consumption; VCO<sub>2</sub>, carbon dioxide production; RER, respiratory exchange ratio; CI, confidence interval for difference in means. n=13 for VO<sub>2</sub> (% peak) measurement from older group.

Carbohydrate and fat oxidation over the 24-hr period are shown in Figure 2. Carbohydrate oxidation was lower and fat oxidation was greater in older compared with young. Carbohydrate oxidation as a percentage of total energy expenditure (Figure 3) was lower, and fat oxidation greater, in the older than the young. On average, there was an energy deficit during the 24-hr stay in both groups, but this did not differ between groups (Young: −1,597 ± 1,612 kJ; Older: −1,313 ± 945; p = 0.539). After adjusting for assumed values of 24-hr protein oxidation from Melanson et al., (2007) and Levadoux et al., (2001), respectively, carbohydrate oxidation remained lower in older (37.4 ± 8.7 kJ·kg^-1^; 39.4 ± 8.7) than young (49.6 ± 12.1, p = 0.002; 50.4 ± 11.6, p = 0.004), while fat oxidation was not different in older (61.1 ± 12.5 kJ·kg^-1^; 64.8 ± 11.6) than young (55.5 ± 12.3, p = 0.205; 57.0 ± 12.5, p = 0.075).

**Figure 2.**
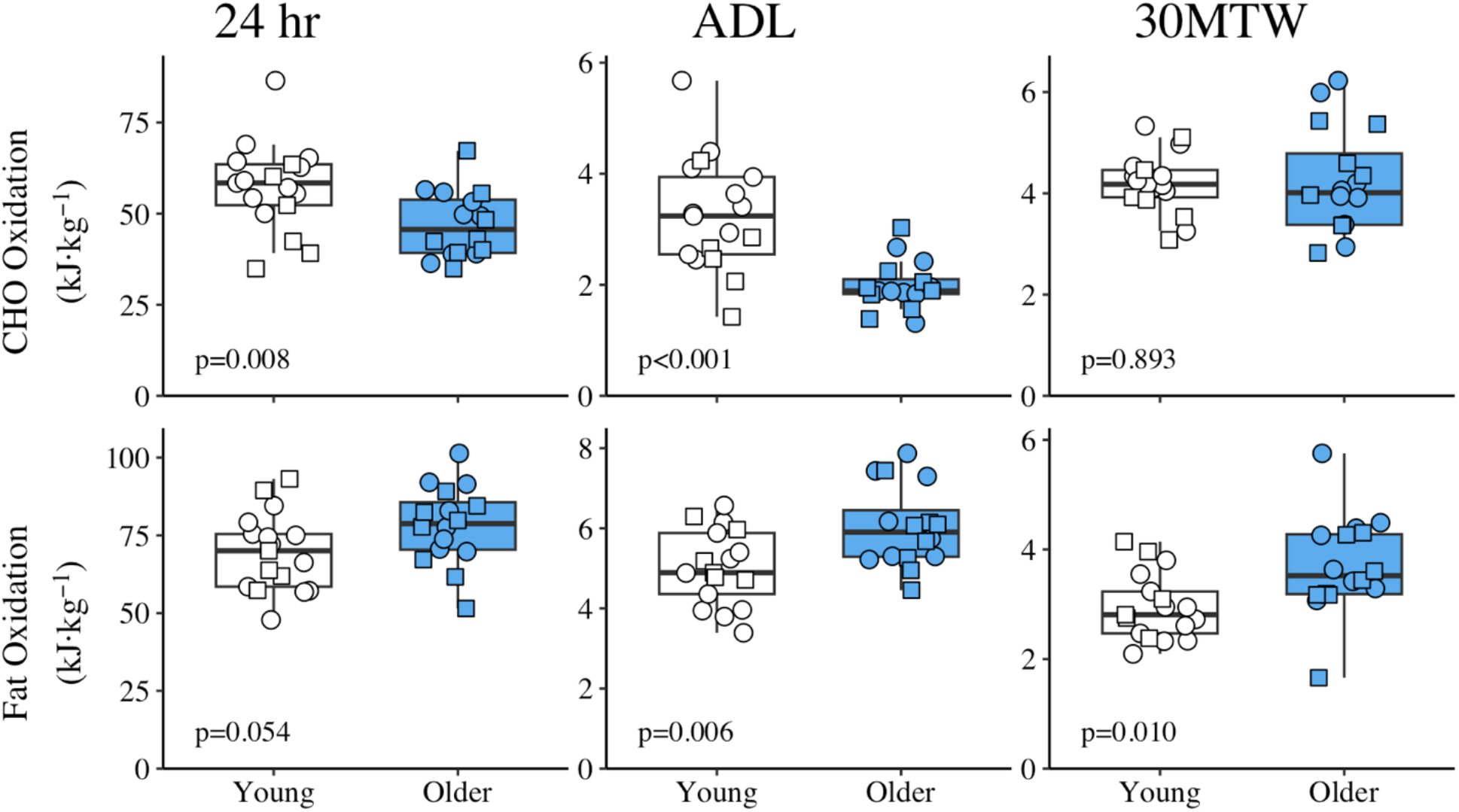
Carbohydrate and fat oxidation for the 24-hour period, activities of daily living, and 30-minute treadmill walk in young and older groups. Carbohydrate oxidation normalized to body mass was lower in older than young over 24 hours (p=0.008) and during ADLs (p<0.001), while body-mass normalized fat oxidation was greater for older than young over 24 hours (p=0.054) as well as during ADLs (p=0.006) and the 30MTW (p=0.010). Carbohydrate and fat oxidation are shown in the top and bottom rows, respectively. Symbols for young adults (n=17) are white and older adults (n=16) are blue. Females are represented by squares and males by circles. Precise p values for between-group differences are provided in the figure for each comparison. ADL, activities of daily living; 30MTW, 30-min treadmill walk.

**Figure 3.**
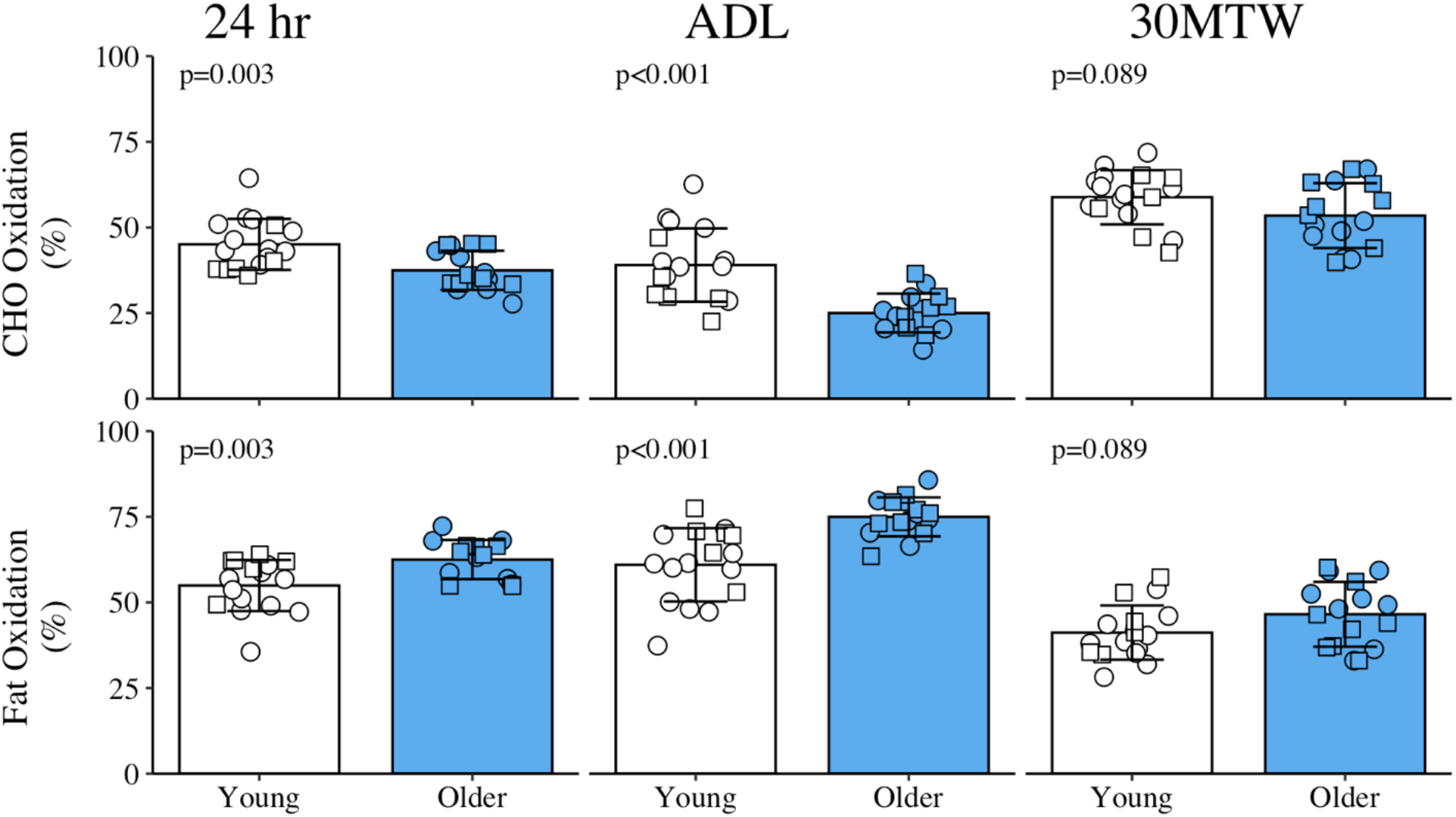
Carbohydrate and fat oxidation relative to total energy expenditure for the 24-hour period, activities of daily living, and 30-minute treadmill walk in young and older groups. The older group oxidized fewer carbohydrates and more fat over 24 hours (p=0.003) and during ADLs (p<0.001) compared with the young group, with no differences in these variables during the 30MTW at 1.3 m·s^-1^ (p=0.089). Carbohydrate and fat oxidation are shown in the top and bottom rows, respectively. Symbols for young adults (n=17) are white and older adults (n=16) are blue. Females are represented by squares and males by circles. Precise p values for between-group differences are provided in the figure for each comparison. ADL, activities of daily living; 30MTW, 30-min treadmill walk.

### Sleep Outcomes

Table 4 shows the data for VO_2_, VCO_2_, energy expenditure, and RER during sleep. Total VO_2_, VCO_2_, and energy expenditure were lower during sleep in the older group compared with the young. When normalized to body mass, energy expenditure and 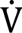O_2_ were not different by age, nor was RER. Oxygen consumption relative to 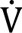O_2peak_ was greater during sleep in the older compared with the young.

**Table 4.** Metabolic variables during sleep.

|  | Young (n=17) | Older (n=16) | p value | 95% CI |
| --- | --- | --- | --- | --- |
| Total VO<sub>2</sub> (L) | 42 ± 7 | 36 ± 7 | 0.015 | 1, 11 |
| VO<sub>2</sub> (ml·kg<sup>-1</sup>·min<sup>-1</sup>) | 3.0 ± 0.4 | 2.8 ± 0.4 | 0.180 | -0.1, 0.5 |
| VO<sub>2</sub> (% peak) | 7.3 ± 0.9 | 10.7 ± 2.1 | <0.001 | -4.8, -2.1 |
| Total VCO<sub>2</sub> (L) | 35 ± 6 | 29 ± 5 | 0.006 | 2, 10 |
| Energy Expenditure (kJ) | 853 ± 150 | 722 ± 134 | 0.012 | 31, 233 |
| Energy Expenditure (kJ·kg<sup>-1</sup>) | 11.0 ± 1.4 | 10.2 ± 1.4 | 0.150 | -0.3, 1.7 |
| RER | 0.82 ± 0.03 | 0.81 ± 0.03 | 0.077 | -0.002, 0.036 |

Data are mean ± SD; VO<sub>2</sub>, oxygen consumption; VCO<sub>2</sub>, carbon dioxide production; RER, respiratory exchange ratio; CI, confidence interval for difference in means. n=13 for VO<sub>2</sub> (% peak) measurement from older group.

Carbohydrate oxidation during sleep was lower in older (3.6 ± 0.9 kJ·kg^-1^) than young (4.6 ± 1.4, p = 0.021), but not when expressed relative to total energy expenditure (Young: 41 ± 9%; Older: 35 ± 9; p = 0.076). Fat oxidation was not different by age when normalized to body mass (Young: 6.4 ± 1.0 kJ·kg^-1^; Older: 6.7 ± 1.5; p = 0.533) or relative to total energy expenditure (Young: 59 ± 9%; Older: 65 ± 9; p = 0.076).

### ADL Outcomes

Mean data for VO_2_, VCO_2_, energy expenditure, and RER obtained during the ADLs are shown in Table 5. Total and body-mass normalized VO_2_ were not different between groups, resulting in the older adults performing this task at a greater percentage of their capacity (% 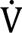O_2peak_). Carbon dioxide production, total energy expenditure, and RER were lower in older compared with young, while energy expenditure normalized to body mass was not different by age. Carbohydrate and fat oxidation during the ADLs are shown in Figure 2. Carbohydrate oxidation was lower and fat oxidation was greater in the older than in the young group. As shown in Figure 3, carbohydrate oxidation as percentage of total energy expenditure was also lower, and fat oxidation greater, in older than young during the ADLs.

**Table 5.** Metabolic variables during activities of daily living.

|  | Young (n=17) | Older (n=16) | p value | 95% CI |
| --- | --- | --- | --- | --- |
| Total VO<sub>2</sub> (L) | 32 ± 7 | 28 ± 5 | 0.070 | -0.3, 8 |
| VO<sub>2</sub> (ml·kg<sup>-1</sup>·min<sup>-1</sup>) | 6.6 ± 0.6 | 6.7 ± 0.8 | 0.926 | -0.5, 0.5 |
| VO<sub>2</sub> (% peak) | 16 ± 3 | 25 ± 5 | <0.001 | -12, -6 |
| Total VCO<sub>2</sub> (L) | 26 ± 6 | 22 ± 4 | 0.017 | 1, 8 |
| Energy Expenditure (kJ) | 655 ± 135 | 562 ± 97 | 0.037 | 5, 166 |
| Energy Expenditure (kJ·kg<sup>-1</sup>) | 8.4 ± 0.8 | 8.0 ± 0.9 | 0.220 | -0.2, 1.0 |
| RER | 0.82 ± 0.03 | 0.78 ± 0.02 | <0.001 | 0.02, 0.06 |

Data are mean ± SD; VO<sub>2</sub>, oxygen consumption; VCO<sub>2</sub>, carbon dioxide production; RER, respiratory exchange ratio; CI, confidence interval for difference in means. n=13 for VO<sub>2</sub> (% peak) measurement from older group.

### 30MTW Outcomes

Oxygen consumption, VCO_2_, energy expenditure, and RER during the 30MTW are given in Table 6. All young and 15 of the older participants were able to maintain the target treadmill speed (1.3 m·s^-1^) throughout the 30MTW. One older female participant walked on the treadmill at a speed of 0.9 m·s^-1^ in order to ensure she could complete the walk. Although total VO_2_ was not different between groups, 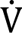O_2_ normalized to body mass and % 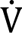O_2peak_ were greater in older than young. Carbon dioxide production during the 30MTW was not different between groups, while RER was lower in older than young. Total energy expenditure was not different between groups, but energy expenditure normalized to body mass was greater in the older group. Carbohydrate and fat oxidation during the 30MTW are shown in Figure 2. Carbohydrate oxidation was not different between groups, while fat oxidation was greater in the older than young. As shown in Figure 3, carbohydrate and fat oxidation as a percentage of total energy expenditure were not different by age.

**Table 6.** Metabolic variables during 30-minute treadmill walk.

|  | Young (n=17) | Older (n=16) | p value | 95% CI |
| --- | --- | --- | --- | --- |
| Total VO<sub>2</sub> (L) | 27 ± 5 | 27 ± 6 | 0.963 | -4, 4 |
| $\dot{V}O_2$ (ml·kg<sup>-1</sup>·min<sup>-1</sup>) | 12 ± 1 | 13 ± 2 | 0.026 | -2, -0.2 |
| $\dot{V}O_2$ (% peak) | 28 ± 5 | 50 ± 13 | <0.001 | -29, -13 |
| Total VCO<sub>2</sub> (L) | 24 ± 5 | 24 ± 5 | 0.826 | -3, 4 |
| Energy Expenditure (kJ) | 558 ± 107 | 558 ± 123 | 0.995 | -82, 82 |
| Energy Expenditure (kJ·kg<sup>-1</sup>) | 7.1 ± 0.4 | 7.9 ± 1.2 | 0.020 | -1.4, -0.1 |
| RER | 0.90 ± 0.03 | 0.87 ± 0.03 | 0.023 | 0.004, 0.05 |

Data are mean ± SD; $\dot{V}O_2$ , oxygen consumption; $\dot{V}CO_2$ , carbon dioxide production; RER, respiratory exchange ratio; CI, confidence interval for difference in means. n=13 for $\dot{V}O_2$ (% peak) measurement from older group.

### Exploring the Crossover Point

By using each individual’s ADL and 30MTW data, which were obtained at different relative exercise intensities, we were able to explore the “crossover” concept for the contribution of fat and carbohydrate oxidation to total energy expenditure in our study groups (Figure 4). As expected, fat oxidation decreased and carbohydrate oxidation increased with greater intensity of exercise, in both groups. Relative to the young group, which had a crossover point at approximately 25% of their 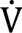O_2peak_, the crossover point for fat and carbohydrate oxidation in the older group appears further to the right (at ∼50% 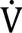O_2peak_), suggesting a lower contribution from carbohydrate to total energy expenditure at the same absolute (Figure 4, *left*) and relative (Figure 4, *right*) intensities.

**Figure 4.**
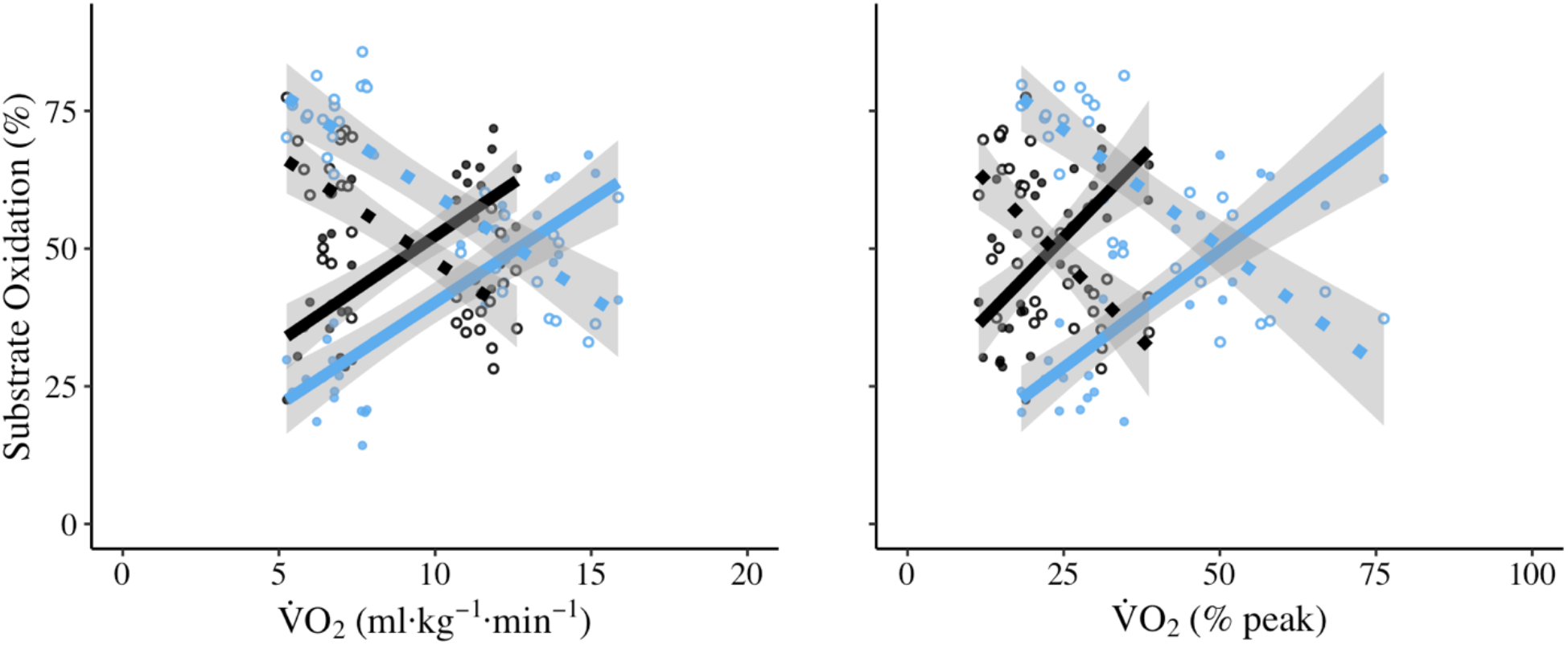
Conceptualization of the “cross-over” point for substrate oxidation in young and older groups as a function of exercise intensity. Plots were generated from individual carbohydrate and fat oxidation data in each group during the activities of daily living and 30MTW, and show the relative contributions from fat and carbohydrate at absolute (*left*) and relative (*right*) intensities. The lines show the fits of the combined within-group data to a linear equation. For both groups, fat oxidation is represented by a dotted line and open circles, and carbohydrate oxidation is represented by a solid line and closed circles. Young adults (n=17) are represented by black symbols and older adults (n=16 on left plot and 13 on right plot) are represented by blue symbols. This preliminary analysis suggests that older adults begin to rely more on carbohydrates for energy at greater absolute and relative intensities compared with young adults.

## DISCUSSION

The goal of this study was to quantify potential differences between young and older adults in energy substrate oxidation over 24 hours and during sub-periods that included sleep, activities of daily living (ADL) and a prolonged period of walking. Our results indicate that carbohydrate oxidation contributed less to the energy requirements over 24 hours (kJ·kg^-1^,%), sleep (kJ·kg^-1^), and ADLs (kJ·kg^-1^,%) of older compared with young adults of comparable habitual physical activity. Conversely, there was a greater contribution from fat oxidation over 24 hours (kJ·kg^-1^, %) and during ADLs (kJ·kg^-1^,%) in older than young adults. Notably, there was no age-related difference in carbohydrate oxidation but an increased reliance on fat oxidation during the 30MTW (kJ·kg^-1^), despite the substantially greater relative intensity during this task in the older group. These results highlight clear age-related differences in substrate oxidation over 24 hours and during daily tasks, which may have important consequences for metabolic health (e.g., metabolic inflexibility, obesity, insulin resistance) and physical function in older age.

### Peak Oxygen Consumption (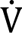O_2peak_)

Our observation of a lower 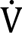O_2peak_ in the older group aligns with the 25-32% lower 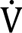O_2peak_ in active older adults reported by Ogawa et al., (1992), who attributed this decline mainly to changes in the cardiovascular system in older age. The lower RER at 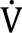O_2peak_ in the older compared with the young group here supports the discussion in the literature (Edvardsen et al., 2014; Wagner et al., 2020) that the widely-used RER criterion of ≥ 1.10 for establishing a maximal 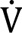O_2_ value may not be appropriate in the older population. However, more work is needed to confirm this concept, and determine the mechanisms for this difference. Regardless, the markedly lower 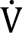O_2peak_ in our older adults clearly demonstrates their lower overall capacity for oxidative energy production, despite maintaining a level of habitual physical activity comparable to that of the young and in excess of the recommended 150 minutes per week of moderate-intensity activity (Allen et al., 2013).

### Diet and Physical Activity

A strength of our study was that we provided a diet while in the room calorimeter based on each participant’s usual diet. These results therefore contribute to our understanding of substrate oxidation in response to a habitual or “native state” diet in these populations, which is likely closer to the metabolic response that may be expected during a normal day. These diets did not differ in macronutrient or caloric content across age groups. Likewise, the time spent lying, sitting, standing or stepping while in the chamber was not different by group. Thus, it is unlikely that our results are related to differences in diet composition or physical activity during the 24-hr room calorimeter stay.

A limitation of this study was the negative energy balance in our groups, likely due to the relatively low caloric intake of the standardized meal bar consumed for lunch. The negative energy balance was nonetheless not different by group, which mitigated any influence of this variable on our between group results. It is possible that performing the ADLs in the fasted state and the 30MTW in an underfed state, following the standardized meal bar, influenced our substrate oxidation results. However, both groups performed these tasks under the same conditions and thus our between-group results are likely not influenced by this to any great extent. We did not measure protein oxidation in this study. In the available literature, protein oxidation over 24 hours is reported to be no different by age (Melanson et al., 2007) or greater in older than young (Davy et al., 2001; Levadoux et al., 2001). However, our analysis using values for protein oxidation in young and older from the literature (Levadoux et al., 2001; Melanson et al., 2007) suggests that including protein oxidation would not have impacted our results or conclusions, particularly the lower 24-hr carbohydrate oxidation in the older than young. Overall, it seems unlikely that these factors influenced the consistent results related to age-group differences in carbohydrate and fat oxidation that we discuss next.

### 24-hr Outcomes

Total energy expenditure over the 24-hour period was not different between young and older adults (Table 3). However, as expected, older adults were at a greater percentage of their aerobic capacity over the day compared with the young group, due mainly to the lower 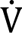O_2peak_ in the older. Despite this greater relative intensity averaged over the day, and contrary to our hypothesis, the older group oxidized fewer carbohydrates and more fat than the young (Figures 2 and 3) over 24 hours. These new results indicate that older adults, compared with young, rely more on fat oxidation for energy production over the course of a day. At present, it is unclear whether the greater use of fat oxidation in this group is a fuel preference, an issue with fuel availability, or a consequence of limited carbohydrate oxidation. Regardless, it is tempting to speculate that this shift toward greater fat oxidation could be related to eventual adverse health outcomes, such as insulin resistance, if it were accompanied by incomplete fatty acid oxidation, which is suggested to have negative downstream consequences on the insulin receptor (Koves et al., 2008). Additional work is needed to identify the effects of greater fatty acid oxidation on metabolic health in aging.

Our results confirm and extend those of Melanson et al., (2007), who found lower carbohydrate (g) and greater fat oxidation (g and g·kg^-1^) in older than young sedentary males over a 24-hour period that included exercise. However, our work contrasts with that of Levadoux et al., (2001) and Davy et al., (2001), who found greater carbohydrate oxidation in older than young and no differences by age, respectively. There are many methodological differences among these previous 24-hour studies and our study that may contribute to the differences we observed. In particular, the lack of objective measures of habitual physical activity between groups in previous room calorimetry studies should be considered, as a more active young group may oxidize more fat and fewer carbohydrates due to training effects, masking any differences by age in substrate oxidation. Our study combined 24-hour substrate oxidation with accelerometry to confirm that our groups were of comparable habitual physical activity levels. Additional potential explanations for the differences between our results and previous work include our relatively larger sample size, recruitment of female participants, use of habitual diet composition, and reporting of units normalized to a metric of body size (kJ·kg^-1^) as well as in both relative (% and RER) and absolute (kJ) terms.

### Sleep Outcomes

Total energy expenditure, but not energy expenditure per body mass, was lower in older than young during sleep (Table 4). The respiratory exchange ratio (Table 4) and substrate oxidation (%) did not differ by age during this period. Our RER results are similar to those of Melanson et al., (2007), who found no difference in RER between young and older males during sleep. However, in contrast to that and other earlier studies (Levadoux et al., 2001), we found that carbohydrate oxidation normalized to body mass (kJ·kg^-1^) was lower in older than young during sleep, indicating that at least a portion of the age-related difference in 24-hr substrate oxidation does not require physical activity in order to be detected in healthy older adults. Further, it appears that the sleep data support the results for the full 24 hours, are consistent with the other sub-periods (discussed below), and highlight the importance of understanding energy metabolism in aging over the full circadian cycle.

### ADL Outcomes

The older group performed the self-paced ADLs at a greater relative (%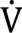O_2peak_) intensity than the young group (Table 5). Notably, energy expenditure per body mass was not different by age during these tasks. Our hypothesis that ADLs would result in greater carbohydrate oxidation in healthy older compared with young adults was not supported by our data (Figures 2 and 3), as we observed markedly lower carbohydrate and greater fat oxidation (kJ·kg^-1^, %), and consequently lower RER in the older adults. These results indicate that the older adults in this study used fewer carbohydrates and more fat, but expended a similar amount of energy (relative to body mass), to perform daily activities at a self-selected pace. It should be noted that the ADLs were performed in a fasted state, which may have influenced substrate oxidation, but would not necessarily explain the difference in substrate use between groups. Nevertheless, these results in combination with the results from the sleep period suggest that the differences by age in 24-hour substrate oxidation were established by metabolic differences at rest as well as during light-intensity physical activities performed during the day.

### 30MTW Outcomes

Oxygen consumption and energy expenditure normalized to body mass were greater for the older than the young during the 30MTW (Table 6), consistent with previous reports of a greater oxygen cost of walking in older age (Hortobágyi et al., 2011; Martin et al., 1992; Waters et al., 1983). Our hypothesis of greater carbohydrate use in older adults during the 30MTW was based on the expectation that they would be operating at a greater percentage of capacity (%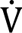O_2peak_) and thus be shifted toward relatively more carbohydrate use during this fixed-speed exercise task. However, this hypothesis was not supported by the data. Although the older walked at a greater relative intensity (%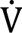O_2peak_) during the 30MTW (Table 6), there was no difference by age in carbohydrate oxidation (kJ·kg^-1^ and %; Figures 2 and 3). Fat oxidation was greater in the older than young (kJ·kg^-1^, Figure 2), though, presumably to meet the increased energy demand of the task and contributing to the lower RER in the older group. Our substrate oxidation results also agree with Hortobágyi et al., (2011), who found a lower RER and greater oxygen cost of walking in older than young while walking at the same absolute workload. Notably, fat oxidation requires more oxygen for a given task compared with carbohydrate oxidation (Krogh and Lindhard, 1920). For example, a high-fat diet has been shown to both increase fat oxidation and oxygen cost in race walkers (Burke et al., 2017). Thus, we propose that the greater oxygen cost of walking in older age may be related to greater fat oxidation in this population. This hypothesis warrants additional investigation.

The results in terms of substrate oxidation during the 30MTW only partially support those observed over the full 24 hours: while RER was lower and fat oxidation greater in the older group, there was no difference in carbohydrate oxidation across groups. It may be that the walking task used here, which was at a greater relative intensity in the older compared with the young group (50% 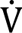O_2peak_ vs. 28%, respectively) was not of sufficient intensity and/or duration to elicit greater carbohydrate oxidation in the young group, but did require more carbohydrate oxidation from the older group. This possibility is supported by the patterns evident in Figure 2 where, relative to the lower-intensity ADLs (Table 5), there was a modest increase in carbohydrate oxidation in the young during the 30MTW, but a near 2-fold increase in the older group. Thus, in combination with their greater fat oxidation (Figure 2), the older group apparently “closed the gap” in carbohydrate oxidation compared with the young in order to meet the energy demands of working at ∼50% 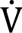O_2peak_. We note that this response suggests that the older group did not have an inability to alter substrate use, and as such were metabolically “flexible” (Goodpaster and Sparks, 2017). Overall, it appears that older adults may need to exercise at a greater relative intensity in order to achieve the same carbohydrate oxidation as young adults. Identifying an appropriate exercise intensity to influence daily substrate oxidation in older adults may be beneficial for metabolic health, particularly as it relates to carbohydrate use and glucose regulation.

### Cross-over Point Concept

Figure 4 places our results from the ADL and 30MTW tasks in the context of the “cross-over” concept of substrate use during exercise (Brooks and Mercier, 1994). Our exploratory analysis shows a cross-over point for the older to the right of the young adults. From this, we estimate that the exercise intensities that elicit substrate oxidation responses of 50% carbohydrate and 50% fat oxidation are ∼50 and ∼25% 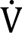O_2peak_ in the older and young, respectively. This conceptual analysis suggests, and supports our points above, that to reach the same substrate oxidation patterns of young adults, older adults may need to perform physical activities at a relatively greater intensity than young. This hypothesis raises a number of questions, including how a “preference” for fat over carbohydrate oxidation might manifest in older muscle, or even whether and how such a shift might be important to metabolic health. Clearly, these results are conceptual, and future work that quantifies the cross-over point on an individual basis during incremental exercise in young and older adults is needed in order to directly address this question and aid in understanding energy metabolism in older humans.

### Potential Mechanisms for Age-related Differences in Substrate Oxidation

There are numerous mechanisms that may contribute to the lower carbohydrate and greater fat oxidation of older compared with young adults found here, including but not limited to age-related differences in substrate availability, product inhibition, enzyme activity, or skeletal muscle fiber type distribution. Regarding the availability of substrates, others have found that [glycogen] is lower in older lifelong endurance athletes compared with young adults (Dubé et al., 2016), and both before and after an endurance training intervention in older compared with young adults (Meredith et al., 1989). Because glycogen availability is positively correlated with glycogen use at submaximal intensities (Hargreaves et al., 1995), lower glycogen availability in the older adults here could have contributed to their lower carbohydrate oxidation during ADLs and over 24 hours. However, reports of greater [glycogen] in older than young (Vigelso et al., 2016), or no difference by age (Nielsen et al., 2010), suggest that this energy substrate may not be limited in healthy older adults. Alternatively, greater fat availability in older age may contribute to our results. Older adults typically have greater intramuscular fat (Dubé et al., 2016; Kent-Braun et al., 2000; Petersen et al., 2015) and circulating free fatty acids (Bonadonna et al., 1994), which may have facilitated greater fat oxidation in the older group found here by providing more substrate. Therefore, although the 24-hour macronutrient content of the diet was not different by group, there is a possibility that substrate availability was different by age and contributed to our results.

The products of greater fat oxidation in older age have been suggested to inhibit carbohydrate oxidation through the Randle cycle (Bonadonna et al., 1994), which maintains that excess acetyl-CoA formation from fat oxidation may inhibit pyruvate dehydrogenase activity and thus reduce carbohydrate oxidation (Hue and Taegtmeyer, 2009; Randle et al., 1963). Evidence of a decline in pyruvate dehydrogenase complex activity in older adults in the insulin-stimulated state (Consitt et al., 2016; Petersen et al., 2015), as well as at baseline and following knee extension exercise (Vigelso et al., 2016), supports this Randle cycle hypothesis. Our results are consistent with the concept that the Randle cycle is amplified in older age; more work is needed to confirm this.

Age-related differences in the activities of glycolytic enzymes should also be considered in the context of our results of lower carbohydrate oxidation in older than young, as these enzyme properties will influence the rate at which carbohydrates can be used. As mentioned above, some researchers have found lower glycolytic enzyme activities in older than young (Consitt et al., 2016; Petersen et al., 2015; Vigelso et al., 2016). However, Coggan et al., (1992) found no differences by age in the activities of multiple enzymes involved in the steps of glycolysis (phosphorylase, phosphofructokinase, and lactate dehydrogenase). Previous work from our lab using magnetic resonance spectroscopy found that glycolysis was not different by age *in vivo* during ischemic muscle contractions (Lanza et al., 2007), suggesting that glycolytic function was maintained in older muscle. Follow-up work is necessary to determine whether age-related differences in glycolytic enzyme activities, or enzymes related to glycolysis (pyruvate dehydrogenase), influence substrate oxidation.

The potential impact of skeletal muscle fiber type on substrate oxidation may be particularly relevant during physical activities. A recent meta-analysis of the literature confirmed early work (Lexell et al., 1988) reporting an age-related fiber-type shift, such that type II muscle fiber area decreases while type I fiber area is not different by age (Lee et al., 2024). As such, type I muscle fibers contribute a greater percentage to muscle volume in older compared with young muscles. Type II muscle fibers generally rely relatively more on glycolysis, and therefore carbohydrate use, than type I muscle fibers (Hultman, 1995). Thus, the lower type II fiber volume in older muscle might explain some of the lower carbohydrate oxidation during exercise in older compared with young adults. The results of Dubé et al., (2016) support this notion, as those authors found that older endurance athletes had fewer type II muscle fibers and lower carbohydrate oxidation compared with young adults while cycling at 40-50% 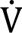O_2peak_.

## Conclusion

The results of this study indicate that, over 24 hours and during sleep and self-paced daily physical tasks, older males and females oxidize less carbohydrate and more fat than young males and females of comparable habitual activity. Carbohydrate oxidation was not different by age, while fat oxidation was greater in older than young, during a 30-min walking task performed at the same speed in both groups, which represented a greater relative intensity for the older adults. This latter finding suggests that older adults met the energy demand of the treadmill walk with relatively more fat oxidation. These results indicate that age-related differences in substrate oxidation are evident during a full day that includes real-world physical activities. The effects of older age on how oxidative energy is derived may be particularly meaningful to understanding the development of adverse metabolic health, such as obesity and insulin resistance, in older adults.

## Data Availability

Data will be made available upon reasonable request to the corresponding author.

## Author Contributions

Conceptualization: ZS, CH, MB, JK. Design: ZS, CH, LA, MB, JK. Funding acquisition: JK, MB. Data acquisition: ZS, CH, LA. Data analysis: ZS, CH, LA, MB, JK. Interpretation: ZS, CH, LA, MB, JK. Drafting and revising the manuscript: ZS, CH, LA, MB, JK. All authors approved the final version of this manuscript and agree to be accountable for all aspects of the work.

## Acknowledgements

We thank the participants of the study. We also thank Joseph Gordon III, PhD, Katie Colfer, Corinna Serviente, PhD, Matthew Limoges, MS, and the undergraduate researchers for their assistance in data collection. We thank Omar Abdelaal for assistance in data analysis.

